# The Influence of Pre-Supplementary Motor Area Targeted High-Definition Transcranial Direct Current Stimulation on Inhibitory Control

**DOI:** 10.1101/2020.10.27.358242

**Authors:** Bambi L. DeLaRosa, Jeffrey S. Spence, Michael A. Motes, Wing To, Sven Vanneste, John Hart, Michael A. Kraut

## Abstract

The neural underpinnings of inhibitory control, an executive cognitive control function, has been a topic of interest for several decades due to both its clinical significance and the maturation of cognitive science disciplines. Behavioral, imaging, and electrophysiological studies suggest that the pre-supplementary motor area (preSMA) serves as a primary hub in a network of regions engaged in inhibition. High-definition transcranial direct current stimulation (HD-tDCS) allows us to modulate neural function to assess cortical contribution to cognitive functioning. The present study targeted HD-tDCS modulation of preSMA to affect inhibition. Participants were randomly assigned to receive 20 min of Sham, Anodal, or Cathodal stimulation prior to completing a semantically cued go/nogo task while electroencephalography (EEG) data were recorded. Both anodal and cathodal stimulation improved inhibitory performance as measured by faster reaction times and increased (greater negative) N2 event-related potentials (ERPs). In contrast, the Sham group did not show such changes. We did not find support for the anodal/cathodal dichotomy for HD neural stimulation. These findings constitute an early investigation into role of the preSMA in inhibitory control and in exploring application of HD-tDCS to the preSMA in order to improve inhibitory control.

## 1 Introduction

Inhibitory control is an executive function necessary for flexible and adaptive behavior. Optimal functioning of brain regions subserving inhibition is important in navigating an often changing environment. Research on inhibitory control has been the focus of many investigations due to the clinical significance and to the theoretical development of formal ontologies of cognitive control mechanisms. Towards both these aims, delineating the brain regions primarily responsible for inhibition as well as investigating neuromodulation techniques to exploit optimal functioning of these regions is of current significance.

### 1.1 Inhibitory control and the neural correlates

Traditionally, the Go/NoGo (GNG) task and the Stop-Signal task (SST) have been two of the main experimental paradigms used to probe inhibitory processes. These tasks involve presentation of a target (Go) in which a response is necessary, and presentation of a non-target (NoGo) in which either an initiated response is halted (SST) or withheld (GNG). Initial investigations of inhibition often used the two tasks interchangeably, implying that inhibitory control is a general mechanism necessary for both withholding a motor response (GNG) and for stopping an initiated motor response (SST)[1]. While over time it has become evident that the two tasks have distinct neural bases [2], some of the brain regions involved in both are the pre-supplementary motor area (preSMA), the inferior frontal cortex (IFC), and the subthalamic nucleus (STN) of the basal ganglia [3][4][5]. Meta-analysis of the literature reporting on the two inhibition tasks conclude that the GNG task engages a fronto-parietal network, with emphasis on the role of the preSMA, and the SST task engages a cingulo-opercular network, with emphasis on the right inferior frontal gyrus (rIFG) and the thalamus [2].

The preSMA appears to play an important role in inhibitory control. Swann and colleagues [6]used a combination of fMRI, macrostimulation diffusion tractography, and electrocorticography (ECoG) and concluded that the preSMA mediates inhibitory control. They showed structural connection between preSMA and rIFG, and that stimulation of the preSMA evoked strong local field potentials in rIFG. In addition, during an inhibitory control task gamma and beta frequency-band EEG activity in the preSMA consistently preceded rIFG activity. There is a rich history on neural underpinnings of inhibition, leading to the current view of the preSMA serving as a primary hub in a network of regions engaged in inhibition.

### 1.2 Inhibitory control and electrophysiological correlates

The N2-P3 event-related potentials (ERP) have been shown to relate to inhibitory processes, but the relationships of these waveform components to specific inhibitory functions remain to be fully specified [7]. The N2 is a fronto-central surface negative polarity waveform component that peaks around 200 ms after stimulus onset, and the P3 is a frontocentral positive peak around 300 ms post stimulus presentation [8][9][10]. It has been postulated that N2 reflects cognitive inhibition or response conflict, while the P3 reflects inhibition of an overt response [11][12]. Both the N2 and P3 peaks are greater for NoGo stimuli than they are for Go stimuli and are delayed with increasing task difficulty.

The GNG task is ideal for investigating the cortical contributions to inhibition, as the task consistently elicits activity in the preSMA [2]. We have previously used a semantically cued GNG inhibition task to probe electrophysiological correlates of inhibitory control [13][14][15][16]. This task requires a semantic categorization assessment of each stimulus before determining to respond (Go) or to withhold a response (NoGo), with different levels of difficulty based on the semantic cue (single object decision or categorical decision). Performance of this task evokes the expected N2-P3 response in frontal midline electrodes, with greater N2-P3 amplitudes for the NoGo trials compared to the Go trials. In addition, the semantically cued GNG task is sensitive to deeper semantic processing, with delayed latencies and attenuated P3 amplitudes for the more difficult level of the task [14]. This semantically cued GNG task has been used as a marker of inhibitory control in healthy adults and is sensitive to changes in inhibitory control via training [16], impairments in Gulf War veterans [17], and developmental changes [15].

### 1.3 Inhibitory control and neuromodulation

Over the past several years, non-invasive transcranial electrical stimulation techniques that modulate neural function have become available to investigate the roles that particular brain regions play in a variety of cognitive functions [18][19][20]. Transcranial direct current stimulation (tDCS) is one of these techniques. TDCS combined with behavioral and electrophysiological measures can be used to study the neural underpinnings of cognitive functioning, such as inhibitory control, and may have therapeutic potential as well. Transcranial-DCS delivers a small current (typically 0.5-2.0 mA) through the surface of scalp via either saline-soaked sponge electrodes or Ag/Ag-Cl EEG electrodes. It has been found that positive polarization (anodal) stimulation excites the underlying cortex, while negative polarization (cathodal) dampens or reduces the likelihood of neuronal firing in underlying tissue [21][22][23][24][25]. To date, few studies have utilized tDCS in modulating response inhibition, and of these there has been a focus on the rIFG during the SST [26][27][28][29], rather than the on the preSMA [30]. Hsu and colleagues delivered tDCS to the preSMA region and showed that anodal stimulation improved inhibition, and that cathodal stimulation impaired inhibition, as evidenced by changes in behavior during a SST task. This was a first step in better understanding the contributions of preSMA in inhibition. However, because traditional tDCS often stimulates a large cortical area it is difficult to rule out contributions from stimulation on surrounding areas.

In order to focus to the greatest extent possible stimulation of the preSMA, we used high definition tDCS (HD-tDCS), which uses smaller electrodes than are typically employed in standard tDCS, to study the effects on electrophysiological and behavioral measures of inhibition during a semantically cued GNG task. We hypothesized that anodal stimulation of the preSMA would improve inhibitory functioning, that cathodal stimulation would impair it, and that ERP markers of inhibition, N2/P3, would show changes that reflect the observed behavioral effects.

## 2 Methods

### 2.1 Participants

Data of forty subjects (19 female; 21 male) were analyzed in this study. Five participants were excluded due to unusable EEG data. Participants were between the ages of 18 and 35 (M = 23.77, SD = 4.19). All participants were right handed, reported no neurological impairments, and gave informed consent prior to participation in accordance with the Institutional Review Board of The University of Texas at Dallas. This study was conducted according to the Good Clinical Practice Guidelines, The Declaration of Helsinki, and the U.S. Code of Federal Regulations.

### 2.2 Stimuli

The GNG task that we used has been described in great detail in previously published research [13][31][32][14][15][16]. The subjects participated in two levels of a semantically cued GNG task. The two levels vary in difficulty based on the level of semantic processing require to respond. For the Single Basic-Level condition (SC), the go stimulus was a single car, and the Nogo stimulus was a single dog. For the Superordinate-Level condition (OA), Go stimuli consisted of drawing of 40 food items, 40 cars, 20 clothing items, 20 kitchen items, 20 body parts, and 20 tools, and the NoGo stimuli consisted of 40 drawings of animals (dogs, lobster, worms, dolphins, etc.).

In each level, there were 160 (80%) ‘Go’ stimuli, for which the subject was instructed to press a button, and 40 (20%) ‘NoGo’ stimuli, for which the subject was instructed to withhold a response. In each level of the task, stimuli were presented for 300ms followed by a fixation point (+) for 1700ms. All of the stimuli were black line drawings fitted to a white 600 x 600 pixel square.

### 2.3 Behavioral Procedures

For each condition there were a total of 200 trials. Instructions for the Single Basic-Level was “to press the response button for a car but not to press the button for a dog.” Instructions for the Superordinate-Level was “to press the response button for all objects but not to press the button for any animals”. Participants were instructed to respond as quickly and accurately as possible. For each condition, there were six versions of the task, in each of which the stimuli were presented in different order; these were counter-balanced across participants and sessions. Participants received a different ordered version for the pre and post stimulation sessions.

### 2.4 HD-tDCS Stimulation

Direct current was transmitted through 5 circular Ag/AgCl Electrodes (1cm radius) with conductive gel on a neoprene head cap and delivered by a battery-driven, wireless multichannel transcranial current stimulator (Starstim tCS®), http://www.neuroelectrics.com). The site for stimulation was determined by the International 10/20 Electroencephalogram System corresponding to FZ for the central electrode and Fp1, Fp2, F7, and F8 for the return electrodes. For the anodal stimulation, the central electrode was programmed as the anode and for cathodal stimulation the central electrode was set as a cathode. The current was initially increased in a ramp-like fashion over several seconds (60 seconds) until it reached 1 mA. The HD-tDCS stimulation procedure was maintained for a total of 20 min and then decreased in a ramp-like fashion over several seconds (60 seconds). For sham HD-tDCS, the placement of the electrodes was identical to real HD-tDCS stimulation. The current in the sham procedure was also increased in a ramp-like fashion over several seconds (60 seconds) until it reached 1 mA. Then, the current intensity was gradually reduced (ramp down) over several seconds (60 seconds) until being switched off. This was followed by 20 minutes without active stimulation.

### 2.5 Behavioral Analysis

Reaction times (ms) were obtained for Go trials for every subject from the onset of each stimulus to the participant’s button push, and trials were rejected if their reaction times were greater than the 99.5th percentile of a fitted gamma function to each subject’s reaction time distribution. A gamma function was used because the reaction time data was right-skewed, rendering standard deviation methods of outlier detection less applicable. Gamma fitting was implemented in Matlab using the gamfit function. No more than 2 trials were discarded per subject per condition. Reaction time values, using correct trials only, were subsequently log transformed and averaged for each subject. Three participants were excluded from behavioral analysis due to a faulty button box (one keypad button not registering all responses).

### 2.6 EEG Recording

Continuous EEG was recorded from a 64-electrode Neuroscan Quickcap using Neuroscan SynAmps2 amplifiers and Scan 4.3.2 software, with a reference electrode located near the calvarial vertex. Data were sampled at 1 kHz with impedances typically below 10 kΩ. Additionally, bipolar electro-oculographic data were recorded from two electrodes to monitor blinks and eye movements (positioned vertically at the supraorbital ridge and lower outer canthus of the left eye). The continuous EEG data were offline high-pass filtered at 0.5 Hz and low-pass filtered at 30 Hz using a finite impulse response (FIR) filter.

### 2.7 EEG Pre-processing

We analyzed the EEG data using scripts developed in our lab that implement functions from EEGLAB version 13.1 [33]running under Matlab 7.11.0. Preprocessing consisted of down-sampling to 512 Hz, removing data recorded from poorly functioning electrodes, and correcting for stereotyped artifacts including eye blinks, lateral eye movements, muscle, line noise, and heart rate using the “Runica” algorithm [33][34], an implementation of the logistic infomax independent component analysis algorithm of Bell and Sejnowski [35]. Stereotyped artifacts were identified by visual inspection of the spatial and temporal representation of the independent components. Continuous data were then segmented into 2-second non-overlapping epochs spanning from 500 ms before to 1500 ms after the presentation of the visual stimuli. Epochs containing high amplitude, high frequency muscle noise, and other irregular artifacts were removed retaining on average 75 percent of all epochs. Finally, missing electrodes were interpolated and data were re-referenced to the average reference [36].

### 2.8 EEG Analysis

ERP’s were calculated for electrodes FZ and CZ [13][14][16][17]for each session (Pre, Post), task level (SC, OA), and condition (Go, NoGo). We performed single-trial baseline correction using the prestimulus interval (-100 ms to 0 ms) as baseline [37]. At electrode FZ the N2 was determined by extracting the largest negative polarity peak within 100-300 ms post stimulus presentation, and the P3 was the largest positive polarity peak within 300-600 ms post stimulus presentation. At electrode CZ the N2 was determined by extracting the largest negative polarity peak within 150-250 ms post stimulus presentation, and the P3 was the largest positive polarity peak within 250-600 ms post stimulus presentation [16].

### 2.9 Statistical Analysis

A mixed-effects linear model with fixed-effects of group (three levels: Sham, Anode, Cathode), session (two levels: Pre, Post), task level (two levels: SC, OA), and condition (two levels: Go, NoGo) was implemented in SAS (Cary, NC) using Proc Mixed. The mixed model also included two random terms to account for subject-level variability and trial-level variability within each subject. The model takes the form *y*_*ijklm*_ = *µ* + *α*_*i*_ + *b*_*j*(*i*)_ + *γ*_*k*_ + *τ*_*l*_ + *δ*_*m*_ + (*αγ*)_*ik*_ + (*ατ*)_*il*_ + (*αδ*)_*im*_ + (*γτ*)_*kl*_ + (*γδ*)_*km*_ + (*αγτ*)_*ikl*_ + (*αγδ*)_*ikm*_ + (*αγτδ*)_*iklm*_ + *e*_*ijklm*_ with *i* as the groups, *j* the number of subjects, *k* is session, *l* is task level, and *m* is condition.

## 3 Results

## 3.1 Behavioral-Accuracy and Reaction Time

For the mixed-effects linear model on accuracy, neither the Group X Session interaction nor higher order Group X Session interaction effects were significant. The main effect of condition (Go, NoGo) was significant, *F*(1,98) = 108.78, *p* <.001. There was lower percent accuracy for the NoGo condition (*M* = 82.90%, *SEM* = 1.09%) compared to the Go condition (*M* = 96.28%, *SEM* = 1.09%), consistent with other research showing more false positives for the NoGo condition, and a higher true positive hit rate for the Go condition.

Results for the mixed-effects linear model on log normalized reaction time for the Go condition revealed a main effect of session (Pre, Post), *F*(1,30) = 13.7, *p* <.001, a main effect of task level (SC, OA), *F*(1,30) = 66.51, *p* = <.001, and an interaction effect of group (Anode, Cathode, Sham) by session by task level, *F*(2,54) = 3.33, *p* =.043. The main effect of session was that participants were faster to respond to Go trials for the post-tDCS session (*M* = 5.68, *SEM* =.02) compared to the pre-tDCS session (*M* = 5.74, *SEM* =.02). The main effect of task level was driven by participants who were faster to respond to the SC level (*M* = 5.62, *SEM* =.02) compared to the OA level (*M* = 5.81, *SEM* =.02). This main effect is a replication of previous studies using this task [14][15]. The interaction effect of group by session by task level was driven by HD-tDCS stimulation. For the OA level the Sham group had no reaction time differences between the post-tDCS session (*M* = 5.82, *SEM* =.05) compared to the pre-tDCS session (*M* = 5.85, *SEM* =.05), *t*(39.8) = −1.15, *p* =.26. For the OA level the Anodal group was faster during the post-tDCS session (*M* = 5.72, *SEM* =.05) compared to the pre-tDCS session (*M* = 5.82, *SEM* =.05), *t*(39.8) = −3.21, *p* =.0026. For the OA level the Cathodal group was faster during the post-tDCS session (*M* = 5.75, *SEM* =.05) compared to the pre-tDCS session (*M* = 5.87, *SEM* =.05), *t*(39.8) = −3.78, *p* <.001. For the Superordinate-Level (OA) there were no changes in reaction time for the Sham group, while both the Anodal and Cathodal stimulation groups had faster reaction times post HD-tDCS stimulation (Figure 1 1). This effect was specific to the Superordinate-Level and was not an overall change in speed, as there was no gain in speed for SC level task performance.

**Figure 1:**
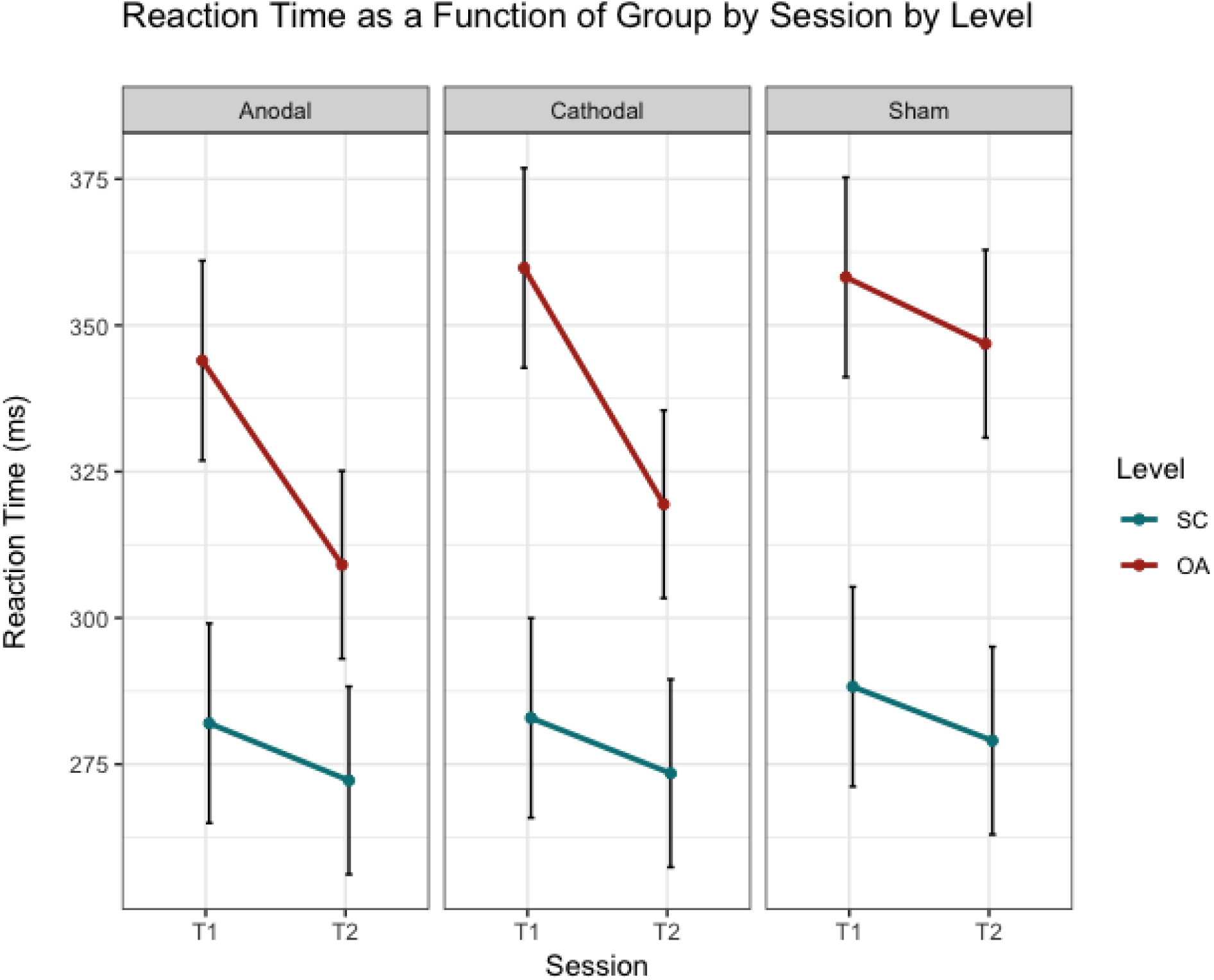
Reaction Time as a Function of Group by Session by Task Level. Reaction time is presented in milliseconds. Both the Anodal and Cathodal group were faster post versus pre for the Superordinate-Level (OA), while there was no change for the Sham group. This faster reaction time was restricted to the OA level as there were no post versus pre stimulation changes for the SC level.

### 3.2 ERP FZ N2

Results for the mixed-effects linear model on N2 amplitudes at electrode FZ replicated previous findings, in that there was a main effect of condition (Go, NoGo), *F*(1,111) = 154.12, *p* <.001, a main effect of task level (SC,OA), *F*(1,111) = 4.29, *p* =.04, and an interaction effect of condition by task level, *F*(1,111) = 7.65, *p* =.006. Pertinent to the aims of this study, there was an interaction effect of group by session by condition by task level, *F*(2,134) = 4.27, *p* =.01. The interaction effect of group by session by condition by task level was driven by HD-tDCS stimulation. For the Superordinate-Level the Sham group had no N2 NoGo amplitude differences at electrode FZ between the post-tDCS session (*M* = −6.76 uV, *SEM* =.70 uV) compared to the pre-tDCS session (*M* = −7.12 uV, *SEM* =.67), *t*(98.3) = 0.65, *p* =.51. For the Superordinate-Level the Anodal group had a greater negative N2 NoGo peak for the post-tDCS session (*M* = −8.07 uV, *SEM* =.65 uV) compared to the pre-tDCS session (*M* = −6.77 uV, *SEM* =.62 uV), *t*(98.3) = −2.50, *p* =.01. For the Superordinate-Level the Cathodal group had a greater negative N2 NoGo peak for the post-tDCS session (*M* = −7.69 uV, *SEM* =.67 uV) compared to the pre-tDCS session (*M* = −6.63 uV, *SEM* =.65 uV), *t*(98.3) = −1.96, *p* =.05. For the Superordinate-Level there were no changes in NoGo N2 amplitude for the Sham group, while both the Anodal and Cathodal stimulation groups had a change in N2 amplitude after HD-tDCS stimulation. This effect was specific to the Superordinate-Level, as there was not a change in N2 amplitude for the Single Basic-Level (Figure 2 2.

**Figure 2:**
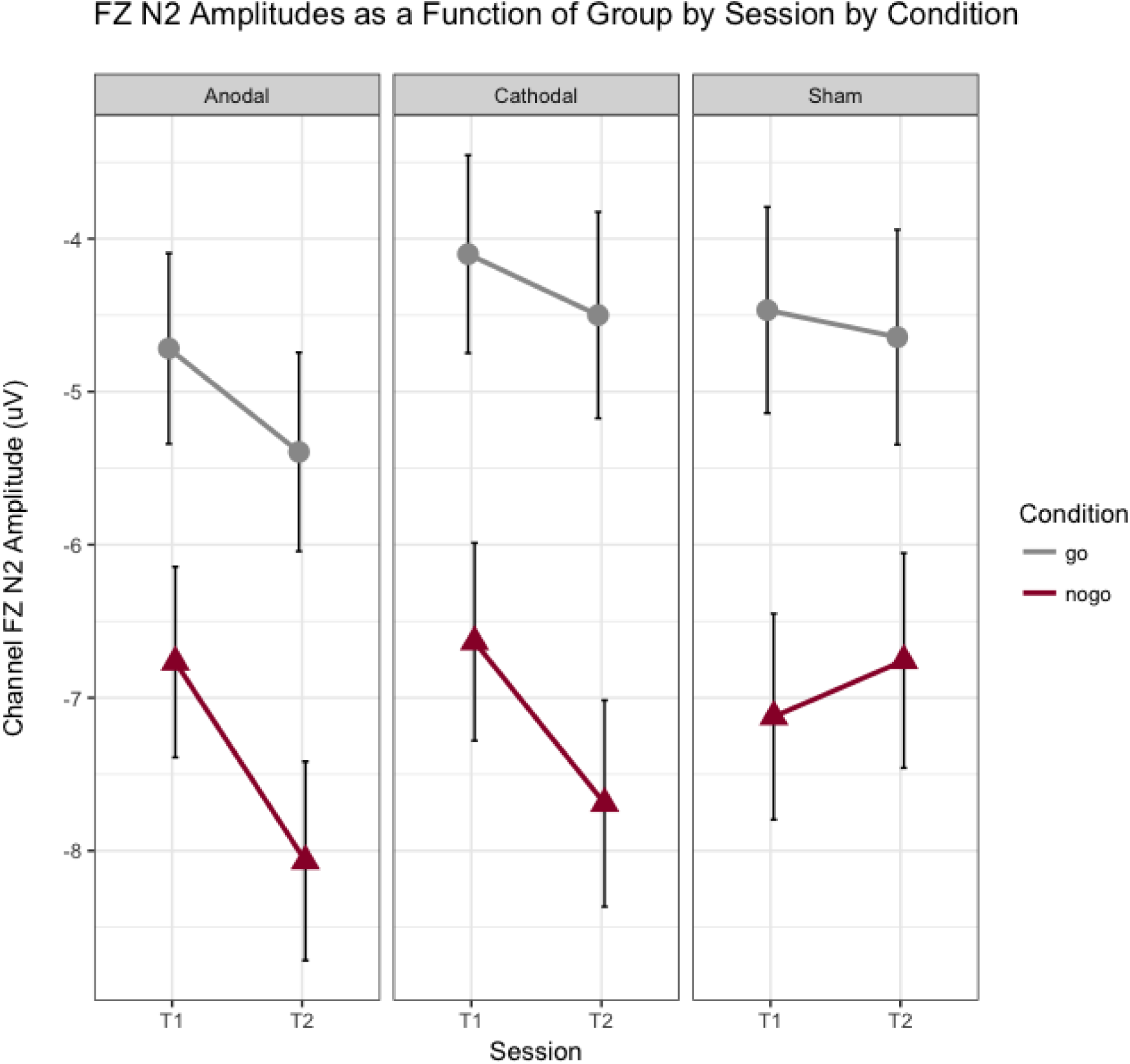
N2 Amplitude at elctrode FZ as a Function of Group by Session by Condition. N2 amplitude is presented in microvolts. Both the Anodal and Cathodal group had greater negative N2 amplitudes for the nogo contition following stimulation (T2), while there were no changes for the Sham group.

### 3.3 ERP FZ P3

Results for the mixed-effects linear model on P3 amplitudes at electrode FZ replicated previous findings, in that there was a main effect of condition (Go, NoGo), *F*(1,112) = 140.31, *p* <.001 and a main effect of task level (SC,OA), *F*(1,112) = 26.94, *p* <.001. There was no effect of stimulation as there was no interaction effect of group by session by condition by task level *F*(2,133) = 2.01, *p* =.14. There was a greater P3 amplitude for the NoGo condition (*M* = 4.43, *SEM* =.21 uV) compared to the Go condition (*M* = 1.93 uV, *SEM* =.21 uV). There was a greater P3 amplitude for the SC level (*M* = 3.73 uV, *SEM* =.21 uV) compared to the OA level (*M* =2.64, *SEM* =.21 uV).

### 3.4 ERP CZ N2

Results for the mixed-effects linear model on N2 amplitudes at electrode CZ replicated previous findings as well was revealed session (pre, post) related attenuation of N2 amplitude. There was a main effect of condition (Go, NoGo), *F*(1,93.3) = 234.42, *p* <.001, an interaction effect of session by task level, *F*(1,140) = 7.68, *p* =.006, and an interaction of condition by task level, *F*(1,93.3) = 4.91, *p* =.03. There was a greater negative N2 amplitude for the NoGo condition (*M* = −5.05 uV, *SEM* =.24 uV) compared to the Go condition (*M* = −2.29 uV, *SEM* =.24 uV). The session by task level interaction effect was driven by N2 attenuation for the SC condition during the post session. The Post SC N2 amplitude (*M* = −3.25 uV, emphSEM =.25 uV) was lower compared to the Pre SC N2 amplitude (*M* = −3.81 uV, *SEM* =.27 uV), *t*(50.8) = 2.54, *p* =.01. There was no session difference for the Post OA N2 amplitude (*M* = −3.76 uV, *SEM* =.25 uV) compared to the Pre OA N2 amplitude (*M* = −3.85 uV, *SEM* =.27 uV), *t*(50.8) =.41, *p* =.68.

### 3.5 ERP CZ P3

Results for the mixed-effects linear model on P3 amplitudes at electrode CZ replicated previous findings as there was a main effect of condition (Go, NoGo), *F*(1,94.8) = 104.84, *p* <.001, and there was a main effect of task level (SC, OA), *F*(1,94.8) = 24.96, *p* <.001. There was a greater P3 amplitude at electrode CZ for the NoGo condition (*M* = 4.69 uV, *SEM* =.19 uV) compared to the Go condition (*M* = 2.76 uV, *SEM* =.19 uV). Also, there was a greater P3 amplitude at electrode CZ for the SC level (*M* = 4.19 uV, *SEM* =.19 uV) compared to the OA level (*M* = 3.26, *SEM* =.19 uV). There were no effects of session or HD-tDCS stimulation at electrode CZ.

## 4 Discussion

Several studies have implicated the preSMA in mediating inhibitory control, and that impaired functioning of preSMA, either through cortical neuromodulation [38][30]or through pathophysiology [Attention Deficit Disorder, Schizophrenia, Parkinson’s Disease, Huntingtons Disease][39][40][41][17], results in poor inhibitory control. In this study, we targeted preSMA with HD-tDCS to assess how modulation of this region affects electrophysiological and behavioral measures of inhibition during a semantically cued GNG task. The goals of this study were to clarify further the role of preSMA in inhibitory control, and to begin to establish neuromodulation parameters for optimal inhibitory functioning. We found stimulation effects indicating improved inhibitory performance exemplified by faster reaction times and increased N2 amplitudes post stimulation for both the Anodal and Cathodal group, while the Sham group did not show such changes. The faster reaction times and increase in N2 amplitude we found for the stimulation groups were isolated to the more cognitively demanding level of the task (OA), and effects of stimulation were not found for the simpler task level (SC), indicating the importance of task difficulty in assessing efficacy of neuromodulation. In addition, we replicated previous task related findings often reported for this GNG task at both the stimulation electrode and a nearby electrode (CZ). There were no stimulation effects found at the non-stimulation site (CZ). This indicates that HD-tDCS has the ability to modulate inhibitory control and that effects of modulation are isolated to the region of interest. To date, this is the first study to investigate preSMA targeted HD-tDCS and the subsequent behavioral and electrophysiological effects. These results build on theories implicating the preSMA as a region mediating inhibitory control, and serve as the foundation for further investigation into the use of HD-tDCS in optimizing inhibitory performance.

### 4.1 Behavioral Changes

We found that preSMA targeted HD-tDCS stimulation facilitated faster reaction times for a semantically cued GNG task. The GNG task that we used has two levels of complexity. The Single Basic-Level of the task is equivalent to traditional GNG tasks in that only one type of stimulus denotes a Go trial, in this case a line drawing of a car, and another stimuli denotes a NoGo trial, in this case a line drawing of a dog. Each trial is the same stimulus (car or dog depending on trial condition). Subjects can perform this basic level task by detecting primarily perceptual features (e.g., the dog’s nose; the front wheel of the car), and little in the way of stimulus categorization is needed to determine whether it is a Go or NoGo trial. The more cognitively demanding level of this task is the Superordinate-Level, in that a categorical judgement must be made in order to determine if it is a Go or NoGo trial (whether the stimulus is one of any number of animals or objects included in the stimulus set). Previous studies have found that increased task complexity results in longer reaction times and delay of the N2 peak [42][11]. We found the same pattern of results at the pre-stimulation performances, but showed that HD-tDCS resulted in faster reaction times for the more difficult level of the task after stimulation. Given such fast reaction times and the limited complexity for the Single Basic-Level task, it may be the case that participants were responding as fast as possible and, subsequently, had no room for improvement for the most basic level of the task (ceiling effect). In contrast, the Superordinate-Level is more complex, giving rise to a delay in reaction time providing the potential with electrical stimulation. Seemingly contrary to our findings, other studies using rIFG targeted tDCS showed that anodal stimulation resulted in longer reaction times for the SST [26][27]; although direct comparison is limited due to differing stimulation site, means of stimulation, and task. Only one study has stimulated preSMA during an inhibitory control task (SST) and they did not find any changes in reaction time as a result of the stimulation [30]. We postulate that we found faster reaction times because HD-tDCS delivered more focal stimulation of the preSMA, and that the added task difficulty facilitated the potential for improvement.

### 4.2 ERP Measures

Similar to the behavioral effects, we found greater N2 NoGo amplitudes post stimulation for the more cognitively demanding level of the task. There were no stimulation effects found for the P3. It has been postulated that the N2 is a neural marker of inhibition, while the P3 is more related to the motor aspects of inhibition [11][7][12]. It may be the case that stimulation facilitated making a semantic decision; however, previous experiments that used the semantically cued GNG task found that the P3 changes in relation to increasing semantic complexity [14]. Since we found no changes in P3 we believe it unlikely that stimulation altered the semantic aspects of the task. In addition, an increase in N2 amplitude may indicate better inhibitory functioning. N2 amplitude is smaller in subjects with high error rates compared to those with low error rates [8]. Given that in this task we see post stimulation increases in N2 amplitudes, faster reaction times, and no increases in error rates, we take this to suggest that preSMA targeted HD-tDCS facilitated improved inhibitory control.

### 4.3 HD-tDCS

The mechanisms by which non-invasive transcranial brain stimulation modulates neural activity in underlying brain regions has yet to be fully understood. The current model is that stimulation acts on the resting membrane potential, affecting both sodium and calcium channels and NMDA receptors [43][44][45]. Anodal tDCS has a depolarization effect that lowers activation threshold (excitation), while cathodal has a hyperpolarization effect increasing the threshold for neuronal firing [21][22][46][25]. This model was built on results recorded during stimulation of the primary motor cortex and subsequent stimulation effects on motor evoked potentials [3][24][47][48]. In the current study, given the anodal/cathodal dichotomy, we theorized that the Anodal group would show an increase in performance while the Cathodal group would experience a negative effect of stimulation. However, we found that both the Anodal and Cathodal groups showed positive changes post HD-tDCS stimulation. We theorize that with focal stimulation, the underlying region, in this case the preSMA, is less affected by the direction of current application. More specifically, regional isolation of current delivery may have the same net effect on underlying region, though the mechanism may be different underneath the anode and cathode stimulation site. For example, during stimulation the electric field vector (EF) contains both tangential and radial components relative to the cortex. The radial component has been hypothesized to modulate synaptic efficacy by effecting the soma, while the tangential component is pathway specific, through terminal hyperpolarization/depolarization [49]. This indicates the importance of the morphology of the afferent pathway relative to the EF. It has been shown that due to the symmetrical orientation of synaptic pathways in the primary motor cortex, the tangential EF have no average effect on synaptic efficacy [49]. However, no studies have delineated the precise mechanism of somatic versus terminal hyperpolarization/depolarization in other regions of the cortex, where morphology differs (various types of dense anatomical networks of excitatory and inhibitory neurons). Given these considerations, we cannot provide a precise mechanistic account of neuronal modulation, and it is likely that differences in cortical cytoarchitecture across brain regions will result in different interactions between electrical stimulation and the neural responses that are elicited.

Other factors that likely play a role in the variability of effects observed across brain regions include the amount of current applied and level of impedance, duration of stimulation, initial state of the targeted region, individual anatomical differences, position and size of the electrode, neural circuitry (i.e. inhibition of some neurons may have a paradoxical improvement in behavior rather than the implied assumption that inhibition has a negative effect), or functional stability when neural systems are exogenously perturbed. These points highlight the complexity of the challenge, and thus requirement for more refined models of the mechanisms by which neurostimulation exerts its effects.

In regards to tDCS polarity effect and cognition, a recent meta-analysis on the polarity effects of tDCS in motor and cognitive domains concluded that anodal stimulation often resulted in excitatory effects as evidenced by improvement in motor and cognitive performance; however, cathodal effects were less consistent and rarely resulted in impairment of performance [50]. Studies utilizing HD-tDCS in other cognitive domains have found positive behavioral effects with cathodal stimulation [51][52][53]. In our study, given the positive behavioral effects and changes in N2 amplitude we posit that both anodal and cathodal stimulation may have influenced dopaminergic fluctuations leading to increase in inhibitory performance. In vivo studies looking at the effects of cathodal and anodal tDCS affected extracellular dopamine levels in the rat striatum. Ten minutes of tDCS increased extracellular dopamine levels in the rat basal ganglia for more than 400 minutes post stimulation [54]. In addition, a relationship has been noted between central dopaminergic systems and NoGo N2 measures [55][56]. More specifically, in healthy individuals higher Eye Blink Rates (EBR), a clinical measure of activity of the central dopaminergic system [57][58][59], correlated with larger and more negative N2 amplitudes and higher accuracy of NoGo trials [56]. As mentioned previously, inhibition depends on basal ganglia-prefrontal interactions [60], and patients suffering from basal ganglia disorders show a reduced and delayed NoGo N2 amplitudes [39][55][40].

### 4.4 Conclusion

This is the first study to look at HD-tDCS stimulation effects of the preSMA and subsequent changes in performance and ERP measures of inhibition, N2/P3. These findings begin to address a potential causal role of the preSMA in inhibitory control, and that HD-tDCS of the preSMA may improve inhibitory control. These initial findings serve as rationale for future clinical applications of HD-tDCS in improving inhibitory control, and as primary steps in further clarifying the mechanistic bases of the observed effects.

